# VSPGx: A High-Accuracy Pharmacogenomics Interpretation Software Solution with Automated CPIC Guideline Integration

**DOI:** 10.1101/2025.11.24.690276

**Authors:** Nathan Fortier, Gabe Rudy, Andreas Scherer

## Abstract

Accurate pharmacogenomic genotype determination and interpretation are essential for personalized medicine, yet existing bioinformatics tools face significant limitations in detecting named alleles, maintaining current allele definitions, and providing comprehensive clinical annotations. We present VSPGx, a pharmacogenomics interpretation software solution that identifies diplotypes from next-generation sequencing data and annotates them against Clinical Pharmacogenetics Implementation Consortium (CPIC) and FDA drug recommendations using automated curation of the latest allele definitions.

We benchmarked VSPGx against established tools including Aldy, PharmCAT, and Stargazer using both synthetic datasets and real-world clinical samples. In a comprehensive synthetic benchmark spanning 3,655 CYP2C9 diplotype combinations, VSPGx achieved 99.97% concordance, matching PharmCAT’s performance and substantially outperforming Aldy (93.08%) and Stargazer (27.06%). Clinical validation using 11 TaqMan OpenArray samples demonstrated 88.2% allele concordance and 89.1% phenotype concordance across 110 gene-sample combinations, with all discrepancies attributed to the benchmark data utilizing outdated allele definitions rather than VSPGx errors. Our automated curation process ensures continuous alignment with current CPIC guidelines, addressing a critical gap in existing pharmacogenomic analysis tools. VSPGx provides a robust, clinically-validated solution for pharmacogenomic analysis that combines high-accuracy diplotype calling with up-to-date, evidence-based drug recommendations.

## Introduction

Pharmacogenomics (PGx) is a discipline at the intersection of genetics and pharmacology, that is rapidly becoming a cornerstone of personalized medicine. It aims to reduce adverse drug reactions and enhance treatment efficacy and precision. Aligning individual genetic profiles with pharmacological data is becoming increasingly important in clinical practice, particularly as drug regimens become more complex and as more drugs with variable genetic responses are identified. The Clinical Pharmacogenetics Implementation Consortium (CPIC) provides evidence-based guidelines linking specific gene variants to actionable drug recommendations, supporting accurate detection and interpretation of pharmacogenomic alleles, making it an essential resource for clinical decision-making [1].

Next-generation sequencing (NGS) technologies, including whole genome sequencing (WGS), whole exome sequencing (WES), and targeted gene panels, have become powerful tools for comprehensive pharmacogenomic profiling. However, accurately calling pharmacogenomic variants from NGS data presents unique technical challenges. Many pharmacogenes, such as CYP2D6, exhibit high sequence homology with pseudogenes, leading to ambiguous read mapping and coverage patterns. Additionally, pharmacogenomic variation extends beyond single nucleotide variants (SNVs) to include complex structural variants (SVs) and copy number variations (CNVs), which are difficult to detect reliably using standard variant calling approaches.

While several bioinformatics tools have been developed to address these challenges, existing solutions face significant limitations. Some tools rely solely on small variant calls and cannot incorporate structural variations. Others rely on coverage-based methods that struggle with highly homologous gene regions. Many tools utilize outdated allele definitions that do not conform to current CPIC guidelines, and few commercial platforms offer comprehensive analysis and reporting capabilities necessary for clinical applications.

We have developed VSPGx, a pharmacogenomics genotype detection and interpretation software solution designed to overcome these limitations. VSPGx identifies pharmacogenomic diplotypes from NGS data and annotates them against CPIC and FDA drug recommendations using the most current allele definitions. This paper presents a comprehensive benchmark evaluation of VSPGx, comparing its performance against established tools including Aldy, PharmCAT, and Stargazer using both synthetic datasets and real clinical array samples. Our results demonstrate that VSPGx achieves high accuracy in diplotype calling while maintaining compliance with the latest CPIC guidelines, making it a robust solution for clinical pharmacogenomic analysis.

### Related Work

Numerous publicly available bioinformatics tools have been developed for the detection of pharmacogenomic alleles from NGS data, including Astrolabe [2], PharmCAT [3], Aldy [4], Stargazer [5], Cyrius [6], and PAnno [7]. These tools can be applied to data from targeted gene panels, WES, and WGS workflows. However, tools such as PAnno, Astrolabe, and PharmCAT are limited to calling alleles defined exclusively from small variants, and do not support the analysis of CNVs and complex SVs.

To address this limitation, tools such as Aldy and Stargazer have been developed to detect SVs based on sequencing coverage. These tools call star alleles by considering observed small variants and SVs. However, their reliance on initial read alignments for the detection of pharmacogenomic alleles can lead to challenges in genes like CYP2D6, where sequences are highly similar or indistinguishable from known pseudogenes. This results in ambiguous coverage patterns, impacting the accuracy of the variant calling process. Additionally, a notable limitation of both Aldy and Stargazer is their lack of support for the GRCh38 reference sequence. The CYP2D6 genotyping tool Cyrius was developed to address the shortcomings of existing algorithms caused by the homology between CYP2D6 and CYP2D7 [6]. While this method is capable of reliably detecting star alleles across the full spectrum of variation, it is limited in scope to the CYP2D6 gene and relies on outdated allele definitions that do not conform with current guidelines.

In addition to the publicly available tools listed above, several commercial variant calling and diplotyping tools have been developed. Notably, the DRAGEN Bio-IT platform can perform variant calling and diplotyping for 20 PGx genes, including the detection of CNVs and fusions. [8] However, this platform only provides limited genotyping output based on outdated allele definitions. PacBio has developed another commercial solution for CYP2D6 star allele diplotyping using long-read sequencing data. [9] While these commercial tools are helpful, they are limited in scope to basic genotyping capabilities and cannot provide more in-depth analysis or report generation.

The challenge of accurately assessing the coverage profile of highly homologous pseudogene regions is difficult to overcome due to the limitations inherent in short-read sequencing data. These short reads lack the length necessary to differentiate between pharmacogenes and their corresponding pseudogenes effectively [10]. A breakthrough solution to this challenge is offered by long-read sequencing technology, which enables the clear and unambiguous mapping of sequences to their gene of origin, overcoming interference from pseudogenes, albeit at the cost of a higher test expense. Additionally, long-read sequencing provides valuable phasing information, facilitating the distinction between ambiguous diplotype states. Multiple studies have demonstrated the accuracy of allele calling when leveraging long-read sequencing technology [11] [12].

### VSPGx Variant Detection and Recommendations

We have developed a software solution that identifies pharmacogenomic diplotypes and annotates them against drug recommendations. By default, diplotypes are annotated against CPIC and FDA recommendations, but custom annotations can also be used. This annotation algorithm relies on the following annotation sources:

- **CPIC and FDA Variants**: Variants used to define the PGx alleles in the CPIC database.
- **CPIC and FDA Structural Variants**: Structural variants used to define alleles in CPIC.
- **CPIC and FDA Diplotypes**: Defines diplotypes and their mapping to phenotype and activity scores.
- **CPIC and FDA Gene Drug Pairs**: Gene-drug association information from CPIC.
- **CPIC and FDA Recommendations**: Drug treatment recommendations from CPIC.
- **PGx Drugs**: PGx drug information from CPIC and DrugBank.

The algorithm begins by identifying the named alleles present in the sample based on the allele definitions specified in the CPIC Variants track. For autosomal chromosomes, this process consists of identifying the best matched diplotypes, which consist of a pair of named alleles for each gene. Once diplotypes have been assigned for each gene, the algorithm looks up the associated phenotype and activity score information along with any actionable dosing recommendations from the Diplotypes and Recommendations annotation sources. An example output from this algorithm is shown in Figure 1.

**Figure 1.**
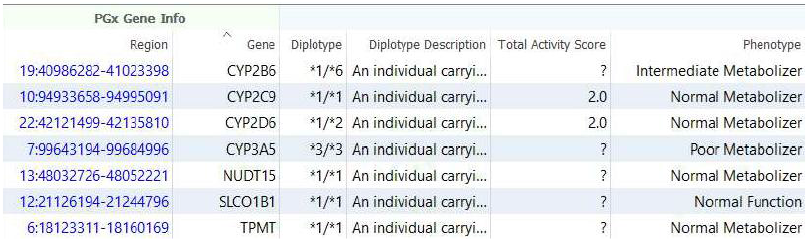
Screenshot of VSPGx Output

Although the algorithm functions on unphased genotype data, it incorporates phasing information whenever available to exclude diplotype configurations incompatible with the observed haplotype structure. Long-read sequencing platforms provide contiguous phasing across the full gene, allowing the algorithm to resolve star-allele ambiguities that remain indistinguishable when using short-read data.

While VSPGx cannot currently detect the presence of SV and CNV alleles, these alleles will be considered by the annotation algorithm if they are specified in the sample table. Similarly, diplotypes called from specialized gene-specific tools like Cyrius can also be incorporated into the analysis, enabling support for a wide range of PGx pipelines.

## Experimental Design

### Synthetic Benchmark

We evaluated the performance of VSPGx’s allele matching algorithm, comparing it to three alternative methods: Aldy, Stargazer, and PharmCAT. This experiment was performed using a simulated benchmark dataset spanning a wide range of CYP2C9 variations. This dataset was constructed by generating all theoretically possible CYP2C9 homozygous and heterozygous diplotype combinations derived from CPIC’s database of named allele definitions. This data was encoded as a collection of 3,655 individual VCF files, each encoding one diplotype state. These files were generated by modifying a reference VCF that starts with all CYP2C9 variants in the default “0/0” state. The steps for generating each file are as follows:

1. Load variants from reference VCF file.
2. For each named allele of the diplotype:
  a. Get the list of core variants defining the allele.
  b. For each variant, update the VCF variant’s genotype to match the alternate allele of the core variant.
3. Save the updated variants to a new VCF.

Using this benchmark dataset, we compared **Aldy, PharmCat, Stargazer**, and **VSPGx** in terms of diplotype, with concordance defined as:

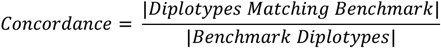

Note that the genotype states stored in the synthetic VCF benchmark files are unphased, making it impossible to distinguish between certain diplotype states. As a result, it is not possible for any diplotype matching algorithm to achieve 100% concordance with the benchmark dataset. However, phasing information could be obtained by leveraging long-read sequencing technology or other supporting assays. When phased genotypes are specified, this information is incorporated into the allele matching process by VSPGx and diplotype states that are inconsistent with the sample’s genotype phasing are excluded from the analysis.

### Clinical Array Sample Benchmark

We obtained data from 11 samples which were analyzed using Applied Biosystems TaqMan OpenArray PGx Express 120 Panel. [13] Each sample was accompanied by an expert-reviewed clinical report that included information on the called alleles, and corresponding drug recommendations based on the patient’s genetic profile.

We converted the provided array data to VCF format, imported the variants and CNVs into VSPGx, and generated clinical reports for each sample using the VSPGx workflow. We compared the original reported alleles and phenotypes to those called by VSPGx and computed concordance for the following CPIC genes: ABCG2, CYP2B6, CYP2C9, CYP2C19, CYP2D6, CYP3A5, DYPD, NUDT15, SLCO1B1, and TPMT.

It is important to note that while these reports offer valuable benchmark data, they were generated based on incomplete and outdated allele definitions. Thus, given VSPGx’s usage of the latest CPIC definitions, perfect concordance was not expected.

## Experimental Results

### Synthetic Benchmark Results

The results of the synthetic benchmark analysis are shown in Table 1.

**Table 1:** Synthetic Benchmark Results.

|  | Aldy | Pharm Cat | Stargazer | VSPGx |
| --- | --- | --- | --- | --- |
| **Concordance** | 93.08% | 99.97% | 27.06% | 99.97% |

PharmCat and VSPGx achieved the best performance, with a concordance of 99.97%. Both algorithms correctly called all but one of the diplotypes in the benchmark dataset. This diplotype was CYP2C9 *10/*12, which was called as *1/*71 by both PharmCat and VSPGx. This mismatch is the result of the ambiguous unphased genotypes in the dataset. The alleles *10 and *12 are defined by the mutations 10598A>G and 50338C>T respectively, while the allele *71 requires the same mutations (10598A>G and 50338C>T) to occur in cis. Thus, it is impossible to distinguish between *10/*12 and *1/*71 without phasing information.

While Aldy has lower concordance than PharmCat and VSPGx, it also performs well, only missing 253 of the 3,655 diplotypes. Stargazer had the poorest performance, with a concordance of just 27%. This low concordance is due to the limitations of the statistical phasing method used by Stargazer, which is ill-suited to rare pharmacogenomic variants. [14] The other three methods do not use a statistical phasing strategy.

These results demonstrate the accuracy of VSPGx’s diplotype matching algorithm. In comparison to other methods, VSPGx achieved a high concordance with the benchmark dataset, correctly calling all but one of the diplotypes.

### Clinical Array Sample Results

The allele and phenotype concordance with TaqMan OpenArray is presented in Table 2.

**Table 2:** Concordance with TaqMan OpenArray.

| Allele Concordance | Phenotype Concordance | Allele Mismatches | Phenotype Mismatches |
| --- | --- | --- | --- |
| 88.2% (97/110) | 89.1% (98/110) | 13 | 12 |

VSPGx achieved high concordance with the alleles and phenotypes included in the clinical reports. Furthermore, all discrepancies in allele calls were traced back to the benchmark reports’ utilization of outdated definitions for six alleles within the CYP2C19 and CYP2D6 genes. **In other words, VSPGx used the most recent CPIC data and correctly classified the alleles, while the benchmark was using outdated information**. There were no discrepancies in the remaining genes where there was agreement on allele definitions.

Through our automated curation process, we ensure that VSPGx always relies on the latest allele definitions from the CPIC database. Consequently, differences between reports generated by VSPGx and those utilizing older allele definitions are not only anticipated but considered favorable, as such differences are a direct result of VSPGx providing recommendations based on the latest CPIC Guidelines.

## Conclusion

This comprehensive benchmark analysis demonstrates that VSPGx achieves high accuracy in pharmacogenomic variant detection and diplotype calling. In our synthetic benchmark evaluation spanning 3,655 CYP2C9 diplotype combinations, VSPGx achieved 99.97% concordance, matching the performance of PharmCAT and substantially outperforming both Aldy (93.08%) and Stargazer (27.06%). The single discordant call in both VSPGx and PharmCAT results from an inherent ambiguity in unphased genotype data rather than algorithmic limitations, highlighting the theoretical limits of diplotype calling from short-read sequencing without phasing information.

Our clinical validation using real-world OpenArray data further confirms VSPGx’s reliability, with 88.2% allele concordance and 89.1% phenotype concordance across 110 gene-sample combinations. Critically, all observed discrepancies were attributed to the benchmark reports utilizing outdated allele definitions, while VSPGx correctly applied current CPIC guidelines. This finding underscores a key advantage of VSPGx: its automated curation process ensures continuous alignment with the latest CPIC database, providing clinicians with recommendations based on the most current evidence-based guidelines rather than legacy definitions.

VSPGx addresses critical gaps in existing pharmacogenomic analysis tools by combining high-accuracy diplotype calling with comprehensive annotation against CPIC and FDA drug recommendations. While the current implementation focuses on small variant detection, the algorithm’s architecture is designed to incorporate structural variant and copy number variation data when specified, positioning it for future expansion as detection methods improve. The platform’s flexible annotation system also supports custom guideline integration, enabling adaptation to institution-specific protocols or emerging pharmacogenomic evidence.

